# Intra-Crater Bubble Expansion Drives the Fracture of Impacted Ureteral Stones in Laser Lithotripsy

**DOI:** 10.1101/2025.06.13.659568

**Authors:** Junqin Chen, Pei Zhong

## Abstract

**Objectives:** To investigate the fracture mechanism of impacted ureteral stones during laser lithotripsy (LL).

**Materials and Methods:** Impacted 6 x 6 mm cylindrical BegoStone samples embedded in a hydrogel ureter model were treated using either Holmium:YAG (Ho:YAG) laser or Thulium Fiber Laser (TFL) via three clinical strategies: “drill and core”, contact, and non-contact modes. Laser pulses were delivered using three pulse energy/frequency settings: 0.8 J/12 Hz, 1.0 J/10 Hz, and 1.2 J/8 Hz with a 3 s on/3 s off protocol, under continuous irrigation at 40 mL/min. Crater formation, surface crack development, and bubble dynamics were assessed via optical coherence tomography, video, and high-speed photography. To delineate the contributions of different plausible damage mechanisms, bubble collapse was suppressed by leveraging the ureteroscope’s proximity effect, and thermal ablation was minimized by treating donut-shaped stones with a central tunnel. The role of bubble expansion in stone fracture was further evaluated by systematically varying tunnel size or pulse energy.

**Results:** Surface cracks and stone fracture were observed exclusively in Ho:YAG laser-treated stones using the “drill and core”, but not the other two strategies. TFL produced deeper craters yet failed to induce any significant crack formation or propagation under all conditions. Suppression of bubble collapse or thermal ablation had minimal effect on surface crack formation and growth. In contrast, the extent and number of surface cracks correlated strongly with the maximum lateral diameter and expansion rate of the vapor bubbles – two parameters that were significantly greater for Ho:YAG laser than TFL. Importantly, the crack formation also showed an inverse correlation with the size of the initial crater or tunnel in the stone.

**Conclusion:** Intra-crater bubble expansion, rather than thermal ablation or bubble collapse, is the primary mechanism driving the fracture of impacted ureteral stone in LL.

## Introduction

Stone dusting and fragmenting represent two distinct treatment modes in laser lithotripsy (LL)^1,2^. While dusting is favored for producing fine debris with shorter procedure time and minimal use of access sheath, fragmenting remains indispensable for managing impacted ureteral or large bladder stones^3–5^. In the past two decades, optimal settings for Holmium:YAG (Ho:YAG) laser have been established, i.e., low pulse energy (E_p_ = 0.2 – 0.4 J) with high frequency (F > 20 Hz) for dusting, and high E_p_ (0.8 – 1.2 J) with low F (< 10 Hz) for fragmenting^3,6,7^. However, for the newer Thulium Fiber Laser (TFL), such parameters are still under active investigations^8–11^. In addition, although TFL offers superior dusting efficiency^12–15^, its ability to fragment impacted calcium phosphate stones in the ureter remains a clinical concern^16,17^. Furthermore, unlike recent progress in elucidating the physical mechanism for stone dusting^8,18,19^, little is known regarding the critical laser parameters and processes that drive stone fracture during LL. Prior studies have not clearly demonstrated whether stone fracture is attributable to either photothermal ablation^20^ or cavitation damage^18,21,22^.

Clinically, dusting is often performed in the kidney using non-contact mode^2,3^, during which stone damage is produced by a combination of photothermal ablation and vapor bubble collapse^8,18–21^. In contrast, fragmenting is preferred for breaking up impacted stones in the ureter into large pieces, followed by basket extraction^1,23^. To avoid ureteral wall injury, the “drill and core” technique has been developed where a small hole is first created at the center of the stone; the fiber is then inserted into the crater to ablate the core while concomitantly fracturing the outer shell^23^. Although widely used, the physical mechanism underpinning this technique for stone fracture has not been explored.

This study aims to elucidate the mechanisms responsible for stone fracture during LL of impacted ureteral stones using either Ho:YAG laser or TFL. Crater damage, surface crack development, and bubble dynamics were compared across three treatment strategies: “drill and core”, contact and non-contact lithotripsy. The mechanism of stone fracture was further investigated using donut-shaped stone phantoms with various central tunnel sizes at different pulse energy levels.

## Materials and Methods

### Impacted stones in a ureter model

To simulate the clinical scenario of impacted ureteral stones, a ureter phantom was constructed using ballistic hydrogel (Gelatin #1, Humimic Medical)^24^ with a 4 mm inner diameter and 8 mm outer diameter, accommodating a snugly fitted 6 x 6 mm cylindrical BegoStone (5:2 powder-to-water ratio, Bego USA) (Fig. 1a). The model was submerged and fixed in a water tank at room temperature (∼20 ^0^C). The flat stone surfaces were treated using a 200 µm core diameter laser fiber (Dornier SingleFlex 200 for Ho:YAG laser and MED-Fibers for TFL, with numerical apertures of 0.26 and 0.22, respectively). The fiber was inserted through the 3.6 F working channel of a digital ureteroscope (Dornier AXIS^TM^) and aligned with the stone center using a 3D translational stage. The scope end was positioned at 5 mm from the stone surface to record the treatment process via its video camera.

**Figure 1.**
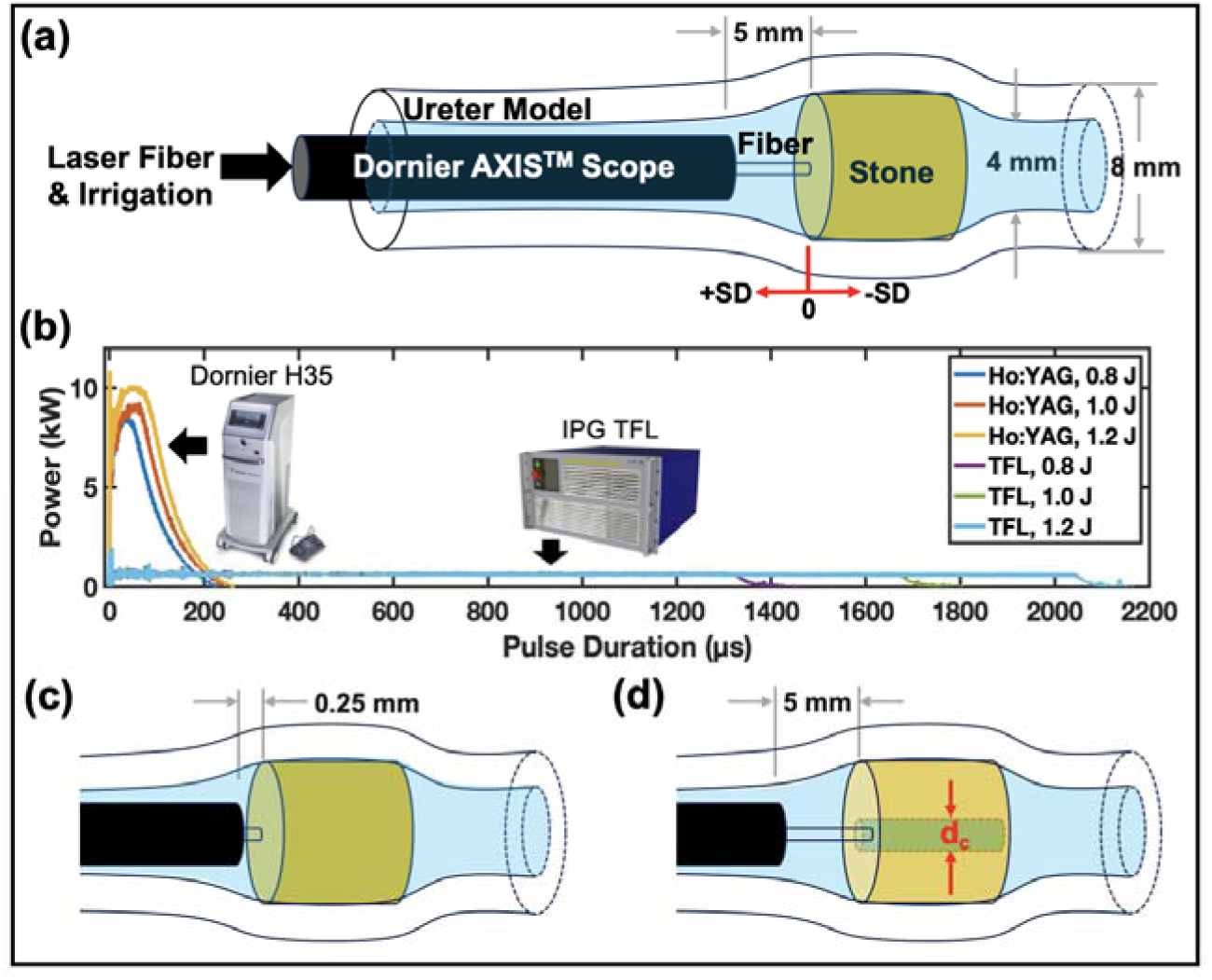
(a) Schematic drawing of a 6 x 6 mm cylindrical BegoStone impacted in a hydrogel-based ureter model and treated by a laser fiber passing through an ureteroscope. The standoff distance (SD) is defined as positive when the fiber is retracted from the stone and negative when inserted into the crater. (b) Pulse profiles of Ho:YAG laser and TFL systems at various pulse energy settings. (c) A cylindrical BegoStone was treated by placing the scope end at 0.25 mm from the stone surface to alter bubble collapse dynamics. (d) A donut-shaped BegoStone with a central tunnel of diameter d_c_ was treated to eliminate the contribution of thermal ablation in front of the fiber tip.

### Laser settings and treatment protocols

A Ho:YAG laser (H Solvo 35, Dornier) and a TFL (TFL-50/500-QCW-AC, IPG Photonics) under three different E_p_/F combinations were used in most of the experiments: 0.8 J/12 Hz, 1.0 J/10 Hz, and 1.2 J/8 Hz. The Ho:YAG laser operated in pre-configured fragmenting mode, while TFL utilized the shortest pulse duration corresponding to a maximum peak power of 500 W (Figure 1b). To prevent excessive temperature rise within the limited fluid volume between the scope end and stone, laser pulses were delivered in 3 seconds on/ 3 seconds off cycles (i.e., 50% duty cycle) under continuous irrigation at 40 mL/min. A total laser energy of 480 J (approximately 48 s lasing time) was delivered, as determined from pilot studies to ensure crack initiation and propagation.

Three treatment strategies were evaluated using pre-soaked BegoStones (n = 15/group):

- ***Strategy 1 (Drill and Core):*** The first 6-second treatment (∼1/8 of the total energy) was delivered at a fiber tip-to-stone standoff distance (SD) of 0 mm, followed by advancing the fiber into the resultant crater at SD = −0.5 mm to complete the treatment, mimicking the clinical “drill and core” protocol^23^.
- ***Strategy 2 (Contact):*** The treatment was performed entirely at SD = 0 mm to maximize photothermal ablation effects^20^.
- ***Strategy 3 (Non-contact):*** The fiber was held at SD = 0.5 mm to enhance bubble collapse effects^22^.

Meanwhile, bubble dynamics were recorded using a high-speed video camera (Phantom v7.3, Vision Research) operating at 50,000 frames per second (fps) either from the side view or at a 45° angle to the laser fiber tip^18^. Additionally, bubble behavior in bulk fluid was recorded using an ultra-high-speed camera (Kirana5M, Specialised Imaging) operating at 200,000 fps^19^.

### Contributions of thermal ablation, bubble collapse and bubble expansion

To further investigate the mechanisms responsible for stone fracture, ***Strategy 1*** was repeated with the ureteroscope end positioned at an offset distance (OSD) of 0.25 mm distal to the fiber tip to reverse the direction of bubble collapse (Fig. 1c)^18^. To mitigate the contribution of thermal ablation in front of the fiber tip and isolate the effect of intra-crater bubble expansion, donut-shaped BegoStones with central tunnels (diameter: 0.49 – 1.17 mm) were fabricated by inserting an optical fiber (diameter: 200 – 1000 µm) into the uncured 5:2 powder-to-water BegoStone slurry to form the tunnel and then removed after 2 hours. After 24 hours of soaking, these donut-shaped samples were treated by advancing the fiber into the tunnel at SD = −0.5 mm (Fig. 1d). All treatments were delivered with a total of 480 J of energy under 40 mL/min continuous irrigation.

### Stone damage quantification and statistical analysis

Post-treatment, crater damage was scanned using optical coherence tomography (OCT; OQ Labscope, Lumedica). Crater volume, maximum depth, and surface profile area were extracted using custom MATLAB code (MathWorks)^22^. Surface crack formation was captured using a high-resolution camera (VMS-005-LCD, Veho), and crack lengths and counts were quantified offline using ImageJ (version 1.54j, National Institutes of Health).

Statistical comparisons across different treatment conditions were performed using Student’s t-tests. Chi-square tests were conducted to assess the influence of laser settings and physical mechanisms on the occurrence rates of distinct damage patterns produced under ***Strategy 1***. A *p*-value < 0.05 was considered statistically significant.

## Results

### Outcomes from different treatment strategies

**Damage Patterns.** Representative results produced at 1.0 J/10 Hz (Fig. 2a) demonstrate distinct differences in stone damage across the three treatment strategies.

**Figure 2.**
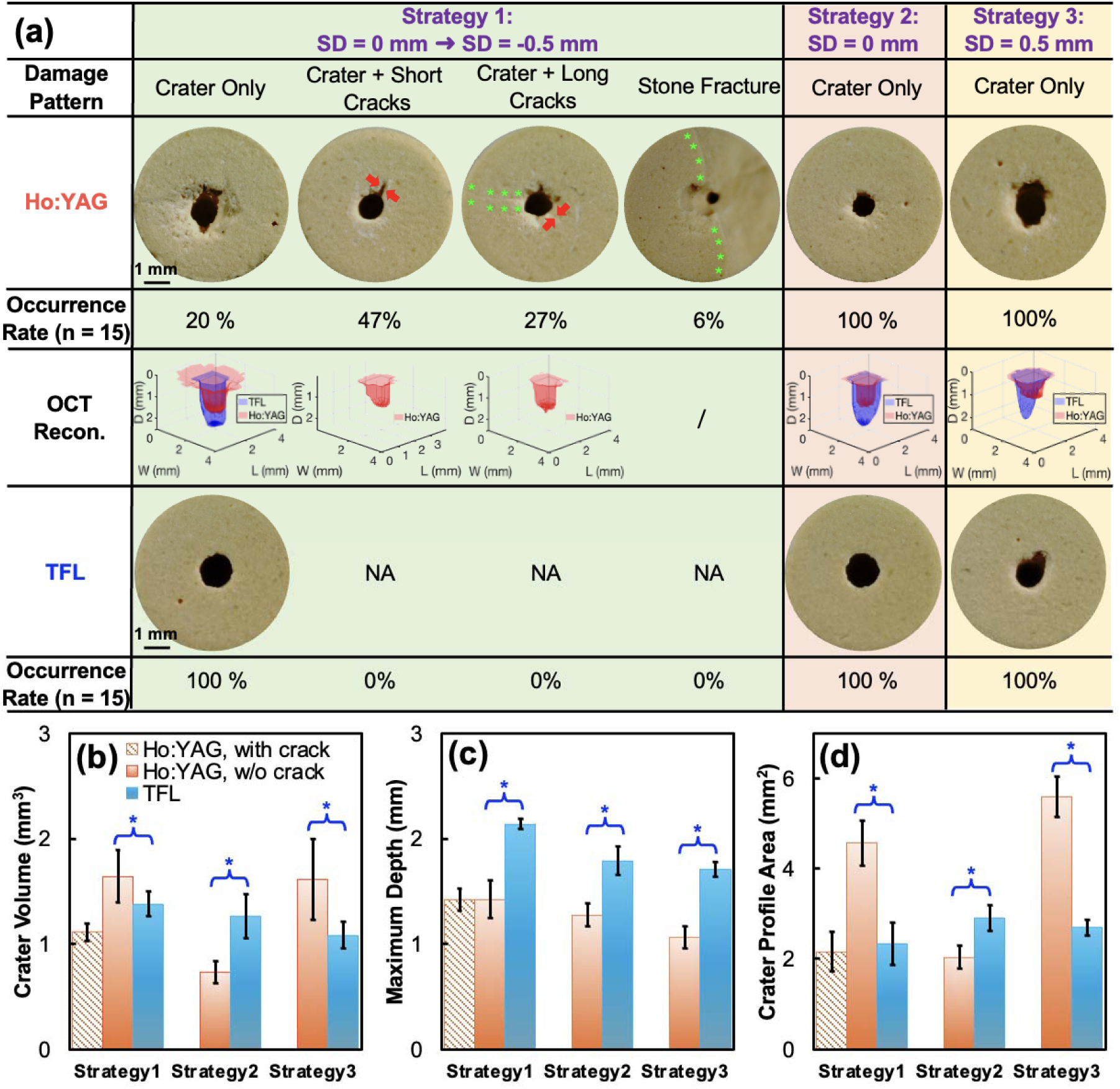
(a) Comparison of representative damage patterns, OCT reconstructions and corresponding occurrence rates across each set of 15 BegoStone samples treated with the Ho:YAG laser and the TFL under different treatment strategies using 1.0 J/10 Hz settings. Red arrows point to the cracks shorter than 1.5 mm (half of the stone radius) and green asterisks denote cracks longer than 1.5 mm. NA = not available. (b) Crater volume, (c) maximum depth, and (d) profile area vs. treatment strategies for the Ho:YAG-treated stones with and without (w/o) cracks, and the TFL-treated stones (all without cracks). All data are presented in mean ± standard deviation with n = 15 for each strategy. *: *p* < 0.05.

Under ***Strategy 1*** (fiber advanced from SD = 0 mm to SD = −0.5 mm), the Ho:YAG laser produced four distinct damage patterns across the 15 treated samples: 1) crater-only (20%), 2) crater with radial cracks shorter than 1.5 mm – approximately half of the stone radius (47%), 3) crater with radial cracks longer than 1.5 mm, extending from the crater edge to the stone boundary (27%), and 4) full stone fracture along radial cracks (6%). Interestingly, among the Ho:YAG-treated stones, the craters without radial cracks had significantly greater volume (*p* = 0.01) than those with cracks, primarily due to their larger profile areas. In comparison, all TFL-treated stones under identical conditions exhibited crater-only damage with no appreciable radial cracks. Nevertheless, the TFL consistently produced deeper craters than the Ho:YAG laser (Fig. 2b-d).

Under ***Strategy 2*** (SD = 0 mm), both lasers produced crater-only damage. However, the craters produced by the Ho:YAG laser were significantly smaller, narrower and shallower compared to the TFL (*p* < 0.001). Yet, no visible radial cracks emanating from the central crater were observed in any of the samples, and the structural integrity of the remaining stone was preserved.

Under ***Strategy 3*** (SD = 0.5 mm), the Ho:YAG laser generated significantly larger crater volumes than the TFL (*p* = 0.007), primarily due to an increase in crater profile areas. Similar to ***Strategy 2***, no radial cracks or stone fractures were observed, and the residual stone remained intact.

**Bubble Dynamics:** Distinct differences in bubble behavior were also observed across the three treatment strategies (Fig. 3).

**Figure 3.**
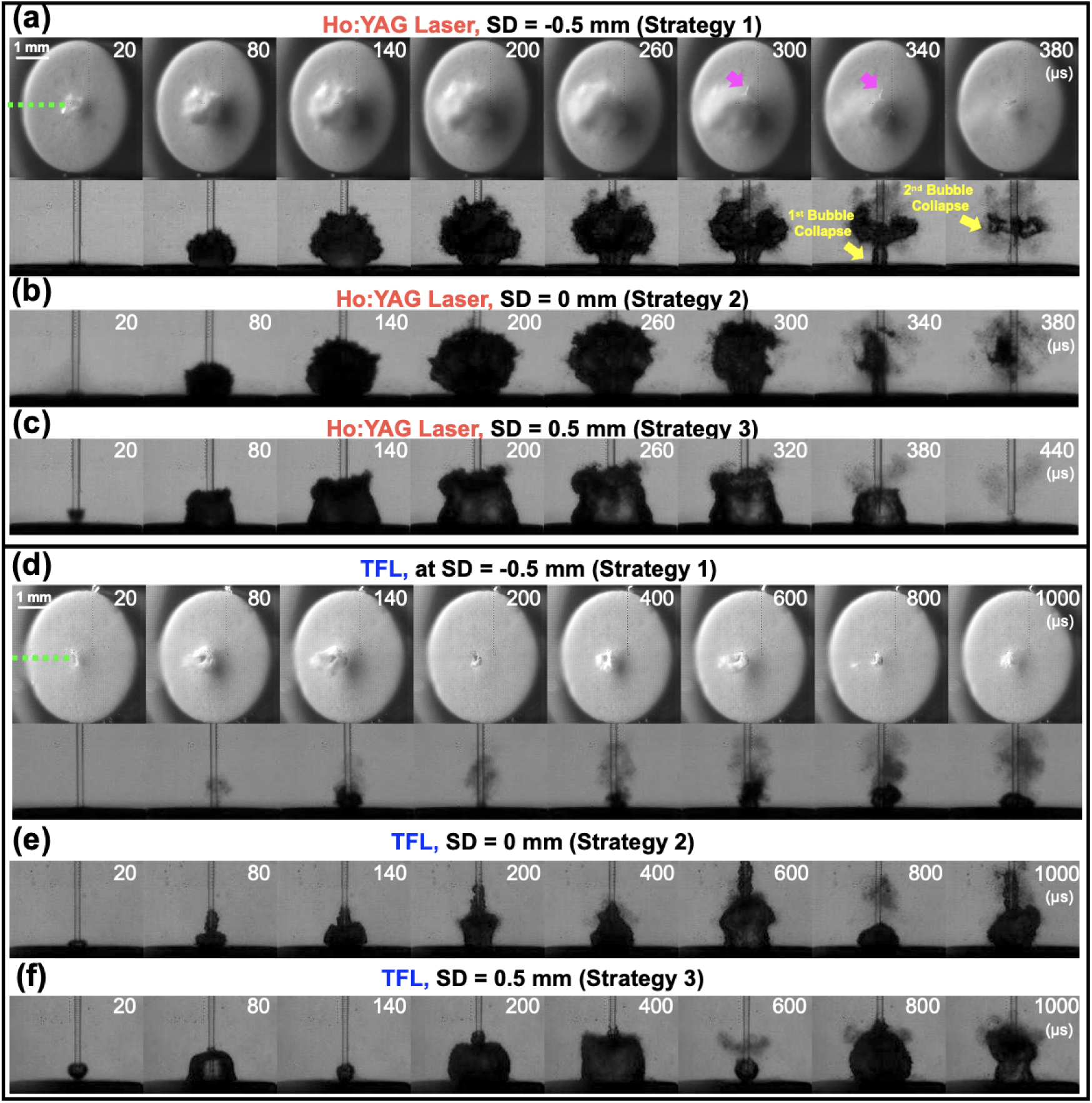
Representative high-speed images of bubble dynamics produced by the 120^th^ pulse of Ho:YAG laser and TFL at 1.0 J/10 Hz settings under (a,d) ***Strategy 1***, (b,e) ***Strategy 2*** and (c,f) ***Strategy 3***. Under ***Strategy 1,*** the interaction between the expanding bubble and surface cracks were captured from a 45° oblique angle. A green dashed line represents the fiber position, pink arrows highlight regions where bubble expansion interacted with surface cracks, and yellow arrows indicate bubble collapse locations from the side view. Additional bubble dynamics produced by different pulse numbers of two lasers are provided in Supplementary Fig. S1 and Fig. S2.

Under ***Strategy 1*** (fiber advanced into the pre-formed crater, SD = −0.5 mm), high-speed images captured at a 45° angle revealed a portion of Ho:YAG laser-induced bubble extending into radial cracks connected to the central crater. This interaction coincided with the onset of crack propagation, first observed at the 120^th^ pulse (see pink arrows of row 1 in Fig. 3a). Side-view imaging showed that the bubble initially expanded within the crater during the first 20 µs, subsequently emerged above the stone surface, and continued to grow irregularly beyond the ∼0.25 mm crater diameter until 260 µs (row 2 Fig. 3a). Two sequential collapse events were captured: the first within the crater at ∼340 µs, followed by a second collapse farther from the stone surface at ∼380 µs. In contrast, under identical settings with TFL, no radial cracks were observed with the 45° angle imaging. Instead, more dust particles were observed dispersing outward (Fig. 3d). No visible bubbles appeared above the stone surface within the first 80 µs. Starting at 140 µs, multiple smaller bubbles emerged and exhibited chaotic expansion and collapse near the fiber tip.

Under ***Strategy 2 (SD = 0 mm)***, the Ho:YAG laser again produced an irregular bubble above the stone surface, though with a larger volume compared to ***Strategy 1*** at the 120^th^ pulse (Fig. 3b). The TFL, on the other hand, generated multiple larger and more irregular bubble oscillations near the fiber tip, accompanied by reduced material ejection compared to ***Strategy 1*** (Fig. 3e).

Under ***Strategy 3 (SD = 0.5 mm)***, the Ho:YAG laser generated a single hemispherical bubble that exhibited the longest expansion-collapse cycle among all three strategies yet resulted in minimal material ejection (Fig. 3c). In comparison, the TFL produced multiple bubbles of different shapes and sizes that were yet overall smaller than those observed with the Ho:YAG laser (Fig. 3f).

**Effect of Crater Size:** Video images captured by the ureteroscope camera demonstrated the effect of the initial crater size on crack formation produced by the Ho:YAG laser under ***Strategy1***. When the initial crater had a small surface profile area (with an equivalent diameter of ∼250 µm; Fig. 4a), fiber insertion into the crater (SD= −0.5 mm) led to the formation of long radial cracks. These cracks began to appear around the 120^th^ pulse and extended to the stone boundary by PN = 180. Additional shorter cracks appeared later (∼ PN = 300), although not all crack events were captured due to partial obstruction of the ureteroscope view by the fiber. Conversely, when the initial crater diameter was large (>600 µm in Fig. 4b), fiber insertion at SD = −0.5 mm did not promote crack formation. Instead, the crater profile area expanded progressively with increasing PN, resulting in crater-only damage.

**Figure 4.**
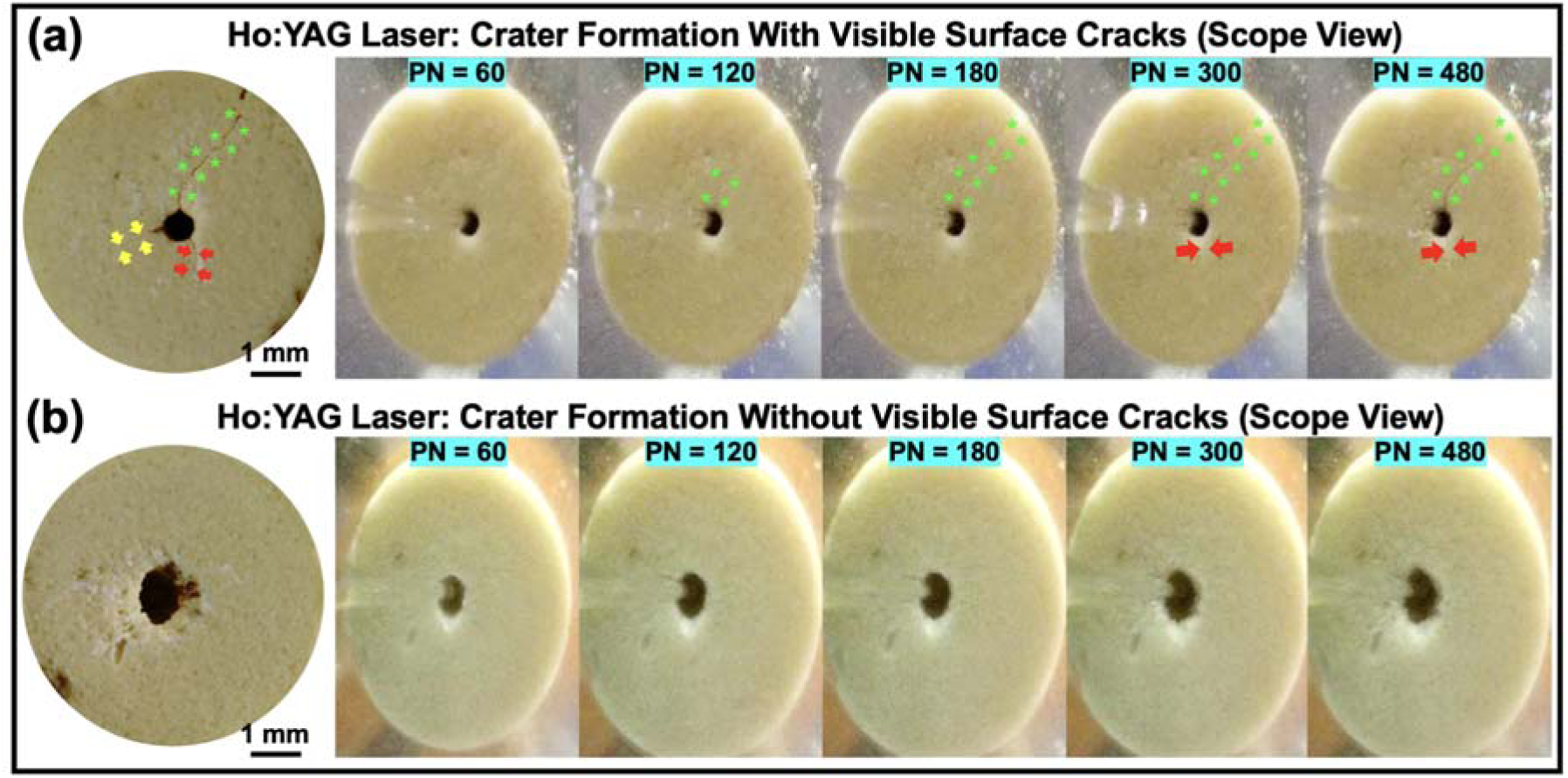
Representative BegoStone samples treated by the Ho:YAG laser at 1.0 J/10 Hz (a) with surface radial cracks propagating from the central crater, and (b) with crater-only damage. Corresponding ureteroscope images at different pulse numbers (PN) are shown on the right. The yellow arrows indicate resultant cracks that were not visible in the ureteroscope view, red arrows point to cracks shorter than 1.5 mm and green asterisks mark cracks longer than 1.5 mm.

### Mechanistic investigation of stone fracture

To further elucidate the underlying physical mechanisms contributing to stone fracture under ***Strategy 1***, we examined the relative contributions of bubble collapse, thermal ablation, and intra-crater bubble expansion.

**Role of Bubble Collapse in Crack Formation:** Bubble collapse was first mitigated by positioning the scope tip at a small OSD of 0.25 mm. Compared to OSD = 5 mm, the bubble produced by the Ho:YAG laser became spatially constrained, exhibiting reduced volume and collapsing toward the scope tip (Fig. 5a-b). Consequently, the crater profile area was significantly reduced (*p* = 0.02). In comparison, no significant change in the crater profile area was observed for the TFL (Fig. 5c). Despite the changes in bubble behavior, surface cracks continued to form with the Ho:YAG laser under both OSD conditions, whereas no cracks were observed in TFL-treated stones (Fig. 5d). Moreover, no statistically significant differences in crack length, number of cracks, or crack occurrence rate were observed between the two OSD cases (*p* > 0.6, Fig. 5d-e), suggesting that bubble collapse plays a minimal role in initiating surface cracks.

**Figure 5.**
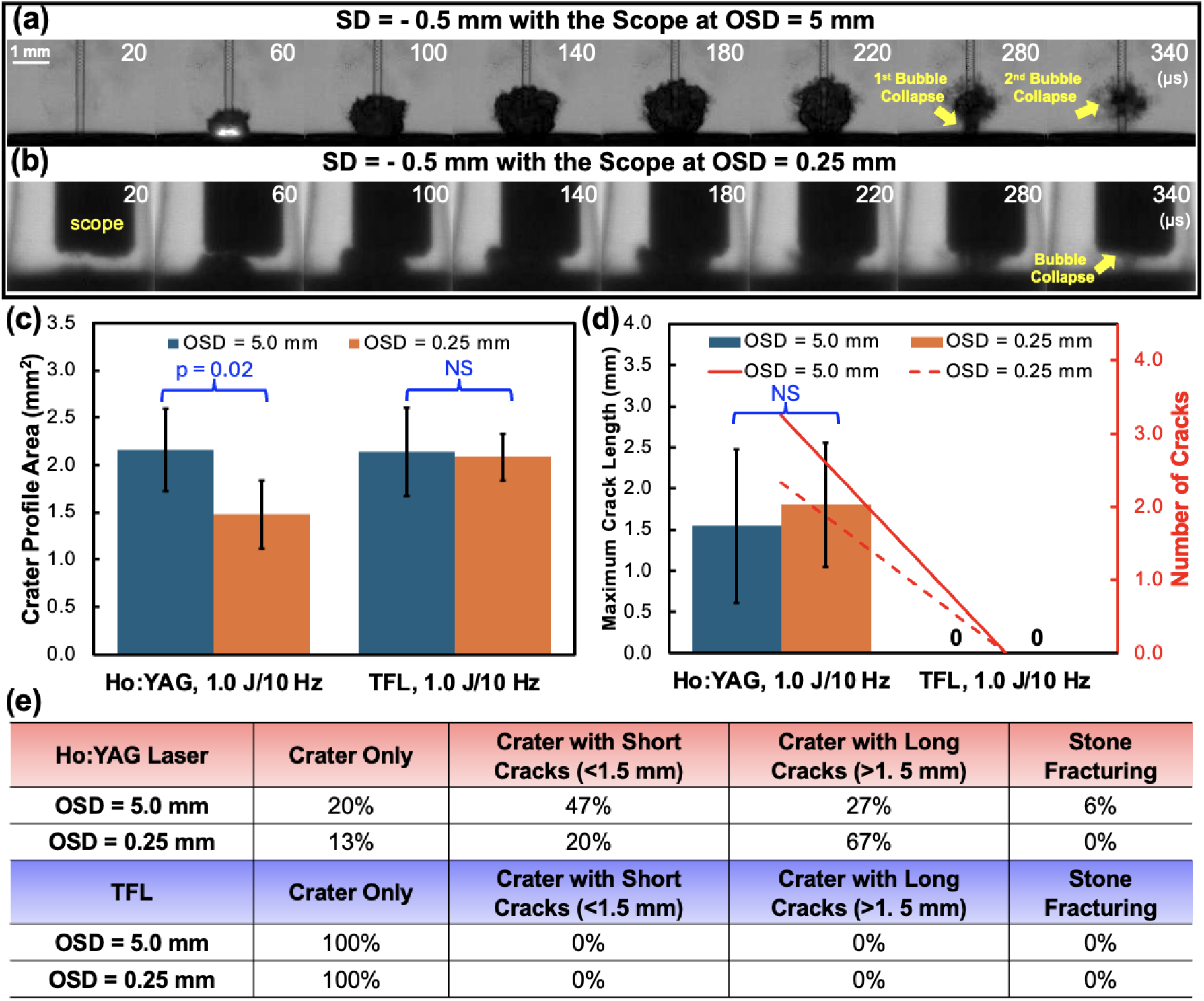
(a) High-speed images of bubble dynamics generated by the 61^st^ pulse of 1.0 J/10 Hz settings with Ho:YAG laser when the fiber tip was advanced into the crater at SD = −0.5 mm and the ureteroscope end was placed at an offset distance (OSD) of (a) 5.0 mm and (b) 0.25 mm. The yellow arrows indicate bubble collapse locations. A comparison of (c) crater profile area and (d) maximum crack length and number of cracks in samples exhibiting craters with short or long cracks, treated by the two lasers at 1.0 J/10 Hz settings under different OSDs. NS: no significance, i.e., *p* > 0.05. (e) The occurrence rates of different damage patterns produced at each OSD, based on 15 samples tested for each group.

**Role of Thermal Ablation in Crack Formation:** The photothermal effect was minimized by treating the donut-shaped BegoStone samples to assess the contribution of photothermal effects to stone fracture. Overall, the cylindrical and donut-shaped stones followed similar trends in the damage outcome across all laser settings (Table 1), suggesting that thermal ablation is not the primary mechanism driving stone fracture. Statistically, for the Ho:YAG laser, a slight reduction in crack length was observed in the donut-shaped samples at 0.8 J (*p* = 0.02). However, Chi-Square analysis showed no significant association between stone geometry and treatment outcomes (p = 0.8). Instead, long radial crack formations were strongly associated with higher E_p_ (*p* = 0.01) regardless of stone shape. In contrast, for the TFL, crater-only damage was consistently observed in both stone shapes across all settings, further supporting that crack formation is not thermally driven in LL treatment.

**Table 1.** Comparison of damage outcomes between cylindrical and donut-shaped BegoStone samples treated with the Ho:YAG laser and TFL across three pulse energy and frequency settings. Data are shown as comparisons between cylindrical and donut-shaped samples. For the crack length and number, values are presented as mean ± standard deviation for the samples exhibiting craters with short and long cracks only.

| Ho:YAG Laser | Crater Only | Crater with Short Cracks (<1.5 mm) | Crater with Long Cracks (>1.5 mm) | Stone Fracture | Crack Length (mm) | Crack Number |
| --- | --- | --- | --- | --- | --- | --- |
| **0.8 J**<br>**10 Hz** | 27% vs. 27% | 60% vs. 67% | 13% vs. 6% | 0% vs. 0% | 0.41±0.25<br>vs.<br>0.21±0.15 | 1.44±0.53<br>vs.<br>1.25 ± 0.12 |
| **1.0 J**<br>**10 Hz** | 20% vs. 13% | 47% vs.54% | 27% vs. 33% | 6% vs. 0% | 1.54±0.93<br>vs.<br>1.29±1.74 | 3.25±1.28<br>vs.<br>3.30 ± 0.61 |
| **1.2 J**<br>**8 Hz** | 13% vs. 13% | 20% vs. 20% | 60% vs. 60% | 6% vs. 6% | 1.93±0.49<br>vs.<br>1.85±0.35 | 3.22±1.30<br>vs.<br>3.25±0.95 |
| TFL | Crater Only | Crater with Short Cracks (<1.5 mm) | Crater with Long Cracks (>1.5 mm) | Stone Fracture | Crack Length (mm) | Crack Number |
| **0.8 J**<br>**10 Hz** | 100% vs. 100% | No cracks observed across all samples. |  |  |  |  |
| **1.0 J**<br>**10 Hz** | 100% vs. 100% |  |  |  |  |  |
| **1.2 J**<br>**8 Hz** | 100% vs. 100% |  |  |  |  |  |

**Role of Bubble Expansion in Crack Formation:** Finally, to examine the role of bubble expansion in stone fracture, two additional control experiments were performed. First, the donut-shaped BegoStones with varying central tunnel diameters (d_c_) were treated by the Ho:YAG laser under 1.0 J/10 Hz settings. Results showed that both crack length and number decreased with increasing d_c_ (Fig. 6a). Second, the donut-shaped stones with the smallest d_c_ of 0.49 mm were treated under different E_p_/F combinations. In this case, crack length and number increased with E_p_ (Fig. 6b). In contrast, the TFL did not produce cracks under any of these conditions. Ho:YAG laser, both the maximum lateral width (*W_b,max_*) and maximum axial length of the To interpret these results, bubble dynamics were further analyzed in free field. For the bubbles increased with E_p_, whereas for the TFL, *W_b,max_* remained nearly constant (∼0.9 mm) and the maximum axial length plateaued at ∼4.0 mm across all energy levels (Supplementary Fig. S3). As a result, the Ho:YAG laser exhibited significantly greater lateral expansion rates 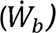 than the TFL while their axial expansion rates were comparable at E_p_ > 1.0 J (Fig. 6c).

**Figure 6.**
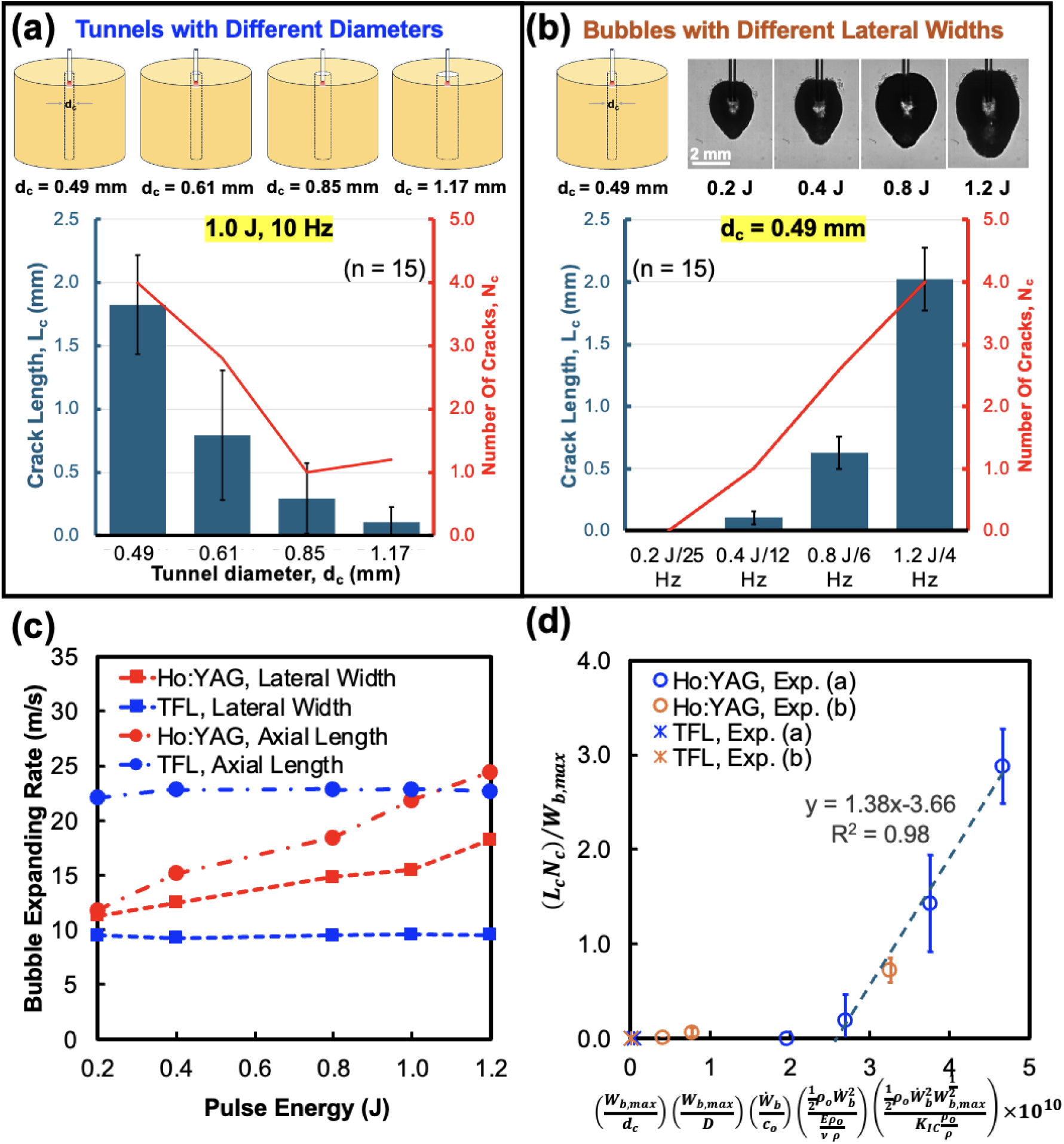
Crack length () and number () in donut-shaped BegoStone samples, which were treated by the Ho:YAG laser and TFL (a) with 1.0 J/10 Hz settings at SD = −0.5 mm and varying tunnel diameter (), and (b) with different pulse energy/frequency combinations by advancing the fiber into the stones with the smallest tunnel (= 0.49 mm) at SD = −0.5 mm. Note: Since the TFL did not induce cracks under all test conditions, its results are omitted from the plots to conserve space. (c) Average lateral and axial expansion rate over the initial 150 µs for Ho:YAG laser-induced bubbles and over the initial 100 µs for TFL-induced bubbles at different energy levels, based on the data shown in Supplementary Fig.S3. (d) The dimensionless group 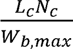, obtained from experimental results, is plotted against 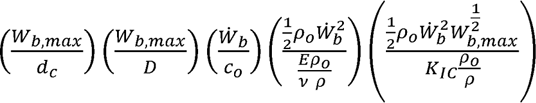 multiplied by 10^10^, where is the maximum bubble lateral width, 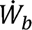 is the average bubble lateral expansion rate, D is the stone diameter, *c*_o_ is sound speed in water, *ρ*_o_ is water density, *E* is Young’s modulus, *v* is Poisson’s ratio, ρ is stone density, and *K*_IC_ is fracture toughness.

Altogether, these findings suggest the critical role of bubble’s lateral expansion in driving crack formation. Through dimensional analysis, we identified two dimensionless groups: 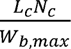, representing the normalized extent of crack formation, and 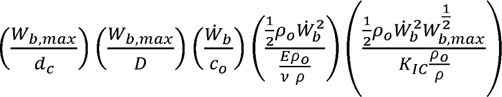, capturing the influence of stone/crater geometry, stone properties and bubble dynamics. Here,*L_c_* is crack length, *N_c_* is number of cracks, D is stone diameter, *ρ*_o_ is water density, *c*_o_ is sound speed in water, *E* is Young’s modulus, *v* is Poisson’s ratio, *ρ* is stone density, and *K*_IC_ is fracture toughness, with values obtained from literature^25,26^. Remarkably, plotting these two dimensionless groups revealed a clear threshold for crack initiation at 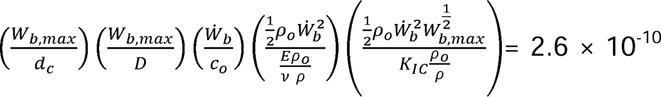, beyond which, the extent of crack formation increased linearly (Fig. 6d).

## Discussion

Stone fragmenting and dusting result in distinctly different treatment outcomes during LL, yet historically, the underlying mechanisms have been attributed broadly to photothermal ablation without clear distinctions^20,27,28^. In this work, we developed a hydrogel-based ureter model to investigate LL of impacted ureteral stones under clinically relevant conditions. Our results demonstrate unequivocally that the Ho:YAG laser can induce stone fracture using the “drill and core” technique (***Strategy 1***), but not by treatment strategies 2 and 3 that only promote either photothermal ablation or stone dusting (Fig. 2a). Notably, the TFL failed to produce stone fractures under any treatment strategies or laser settings tested, despite generating deeper craters than the Ho:YAG laser (Fig. 2c). Mechanistic investigations revealed that stone fracture is less attributable to photothermal ablation or vapor bubble collapse, and more dominantly associated with intra-crater bubble expansion (Figs. 5 and 6, Table 1). Specifically, fracture formation strongly correlates with the maximum lateral expansion size and expansion rate of the vapor bubble, and inversely with the size of the initial crater or central tunnel created at the surface of the stone (Fig. 6). Unlike dusting, which is primarily driven by superficial material removal via photothermal ablation for TFL^8,27^ or vapor bubble collapse for Ho:YAG laser^18,19^, fragmentation requires the initiation of surface cracks that will propagate radially outward from the treatment site to yield bulk fracture (Figs. 2a and 4a).

These novel findings provide a mechanistic explanation for the clinical observation that TFL is less effective than Ho:YAG laser in fragmenting impacted ureteral stones^16,17^. This difference can be attributed to their distinctly different pulse profiles and associated bubble dynamics (Fig. 1b and Fig. S3). The Ho:YAG laser delivers short and shark fin-shaped pulses of high peak power that generate large pear-shaped vapor bubbles characterized by rapid and large lateral expansion. Within the confined geometry of small craters, such bubble expansions can produce strong tensile or shear stresses against the stone boundary, facilitating crack initiation and eventually stone fracture in over 80% of the stones treated by the Ho:YAG laser (Fig. 2a and Fig. 5e). TFL, by contrast, delivers long rectangular pulses with low peak power, which, combined with much higher absorption coefficient in water, produce small yet elongated bubble streams that preferentially expand along the axial direction (Fig. 6c). Consequently, the tensile or shear stresses exerted on the stone wall are presumably much lower than those produced by the Ho:YAG laser, leading to an absence of crack formation under all tested conditions (Fig.2 and Fig. 6).

To further elucidate the origin of mechanical stresses driving stone fracture, we draw parallels to fracture mechanics described in borehole explosion studies^29–31^. Two dominant mechanisms are likely at play here: (1) circumferential tensile (hoop) stress generated along the crater/tunnel wall due to bubble’s lateral expansion, and (2) internal pressurization along crack surfaces due to fluid intrusion. While circumferential tension may produce multiple short, surface-initiated cracks, internal pressurization is more likely to drive fewer but longer cracks, which are consistent with the long radial cracks observed in our experiments (Fig. 3a and 4a). Notably, once two or more long surface cracks forms, continued laser delivery within the crater leads to full stone fracture, as observed in the 6% of Ho:YAG laser-treated stones (Fig. 2a).

The clinical implication of these findings is two-fold. First, for the Ho:YAG laser which inherently promote rapid and large lateral bubble expansion, the initial crater size is critical for successfully initiating stone fracture. Smaller craters enhance confinement and maximize stress generation on the crater wall. As the crater enlarges, increasing the pulse energy may help maintain effective fracture potential by producing larger bubbles. Second, TFL’s low fragmentation potential is intrinsically related to the long bubble stream with small size and limited lateral expansion (Fig. 3, Fig. S2 and Fig. S3). To enhance the fragmentation potential of TFL for impacted ureteral stones, either higher peak power or refined fiber positioning and treatment strategy may be required in the future.

The main limitation of this study is the use of BegoStone phantoms, which do not fully replicate the heterogeneity of human kidney stones, including their layered structure or intrinsic flaws. In addition, bubble dynamics were primarily evaluated in bulk fluid to characterize the maximum expansion size and rate. Future work will incorporate human stones and high-fidelity numerical simulations of the intra-crater bubble-stone interactions^32,33^.

## Conclusion

This study has identified intra-crater bubble expansion as the dominant mechanism for the fracture of impacted ureteral stone in LL using the “drill and core” strategy. While TFL is more conducive to thermal ablation, the Ho:YAG laser offers greater cavitation damage potential, with treatment outcomes influenced by fiber tip positioning and treatment strategy. Unlike dusting produced by thermal ablation or vapor bubble collapse, fracturing of impacted ureteral stones depends critically on surface crack initiation and extension driven by the rapid and large lateral expansion of vapor bubbles produced in a confined crater or tunnel inside the stone. This new discovery clarifies the clinically observed differences in stone fragmentation potential between Ho:YAG laser and TFL. The mechanistic insights gained may help to further refine the treatment strategy to ensure safe and effective fragmentation of impacted stones in the ureter during LL.

## Acknowledgements

The authors are grateful to Dornier *MedTech* for providing the H Solvo laser and the Axis single-use flexible ureteroscope, and IPG Photonics for their technical support with the TFL system. The authors thank Dr. Arpit Mishra and Dr. Obed Isaac for their careful review and valuable feedback on the manuscript. The senior author would also like to express his gratitude to Dr. Margaret Pearle, M.D., Ph.D. of UT Southwestern for recommending the investigation of impacted ureteral stones by TFL at the 2023 ROCK Society Meeting in Boston, MA. This project is supported by the National Institute of Health (NIH) through grants 1P20DK135107-03, 2R01DK052985-27, 1R01DK138972-01A1 and 1R01DK139109-01.

