## Supplementary Fig. S1 and Fig. S2 for "Intra-Crater Bubble Expansion Drives the Fracture of Impacted Ureteral Stones in Laser Lithotripsy"

**Supplemental Data**

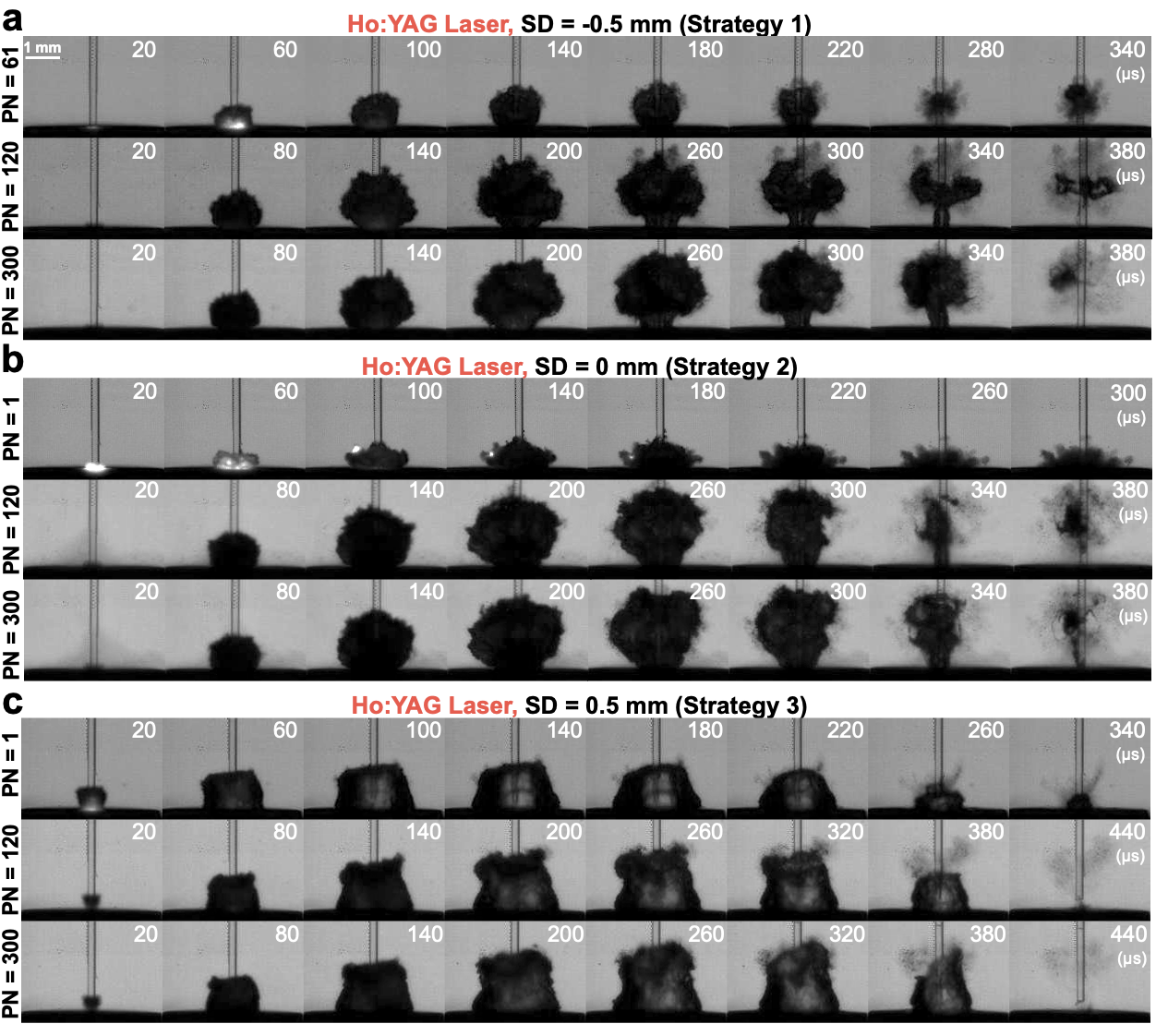

**Figure S1.** Representative high-speed images of bubble dynamics produced by different pulse number (PN) of Ho:YAG laser at 1.0 J/10 Hz settings under (a) ***Strategy 1***, (b) ***Strategy 2*** and (c) ***Strategy 3***.

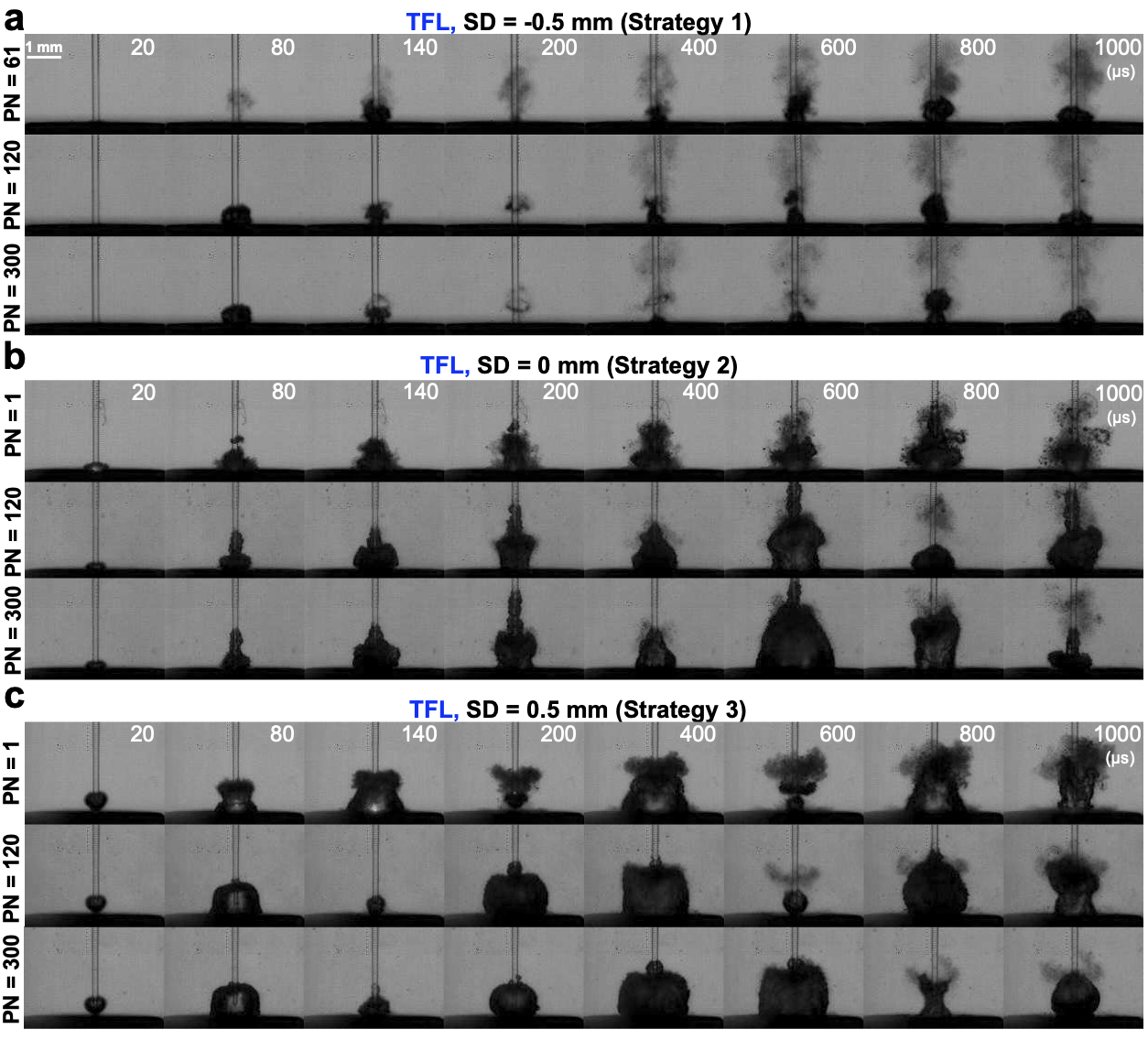

**Figure S2.** Representative high-speed images of bubble dynamics produced by different pulse number (PN) of TFL at 1.0 J/10 Hz settings under (a) ***Strategy 1***, (b) ***Strategy 2*** and (c) ***Strategy*** ***3***.

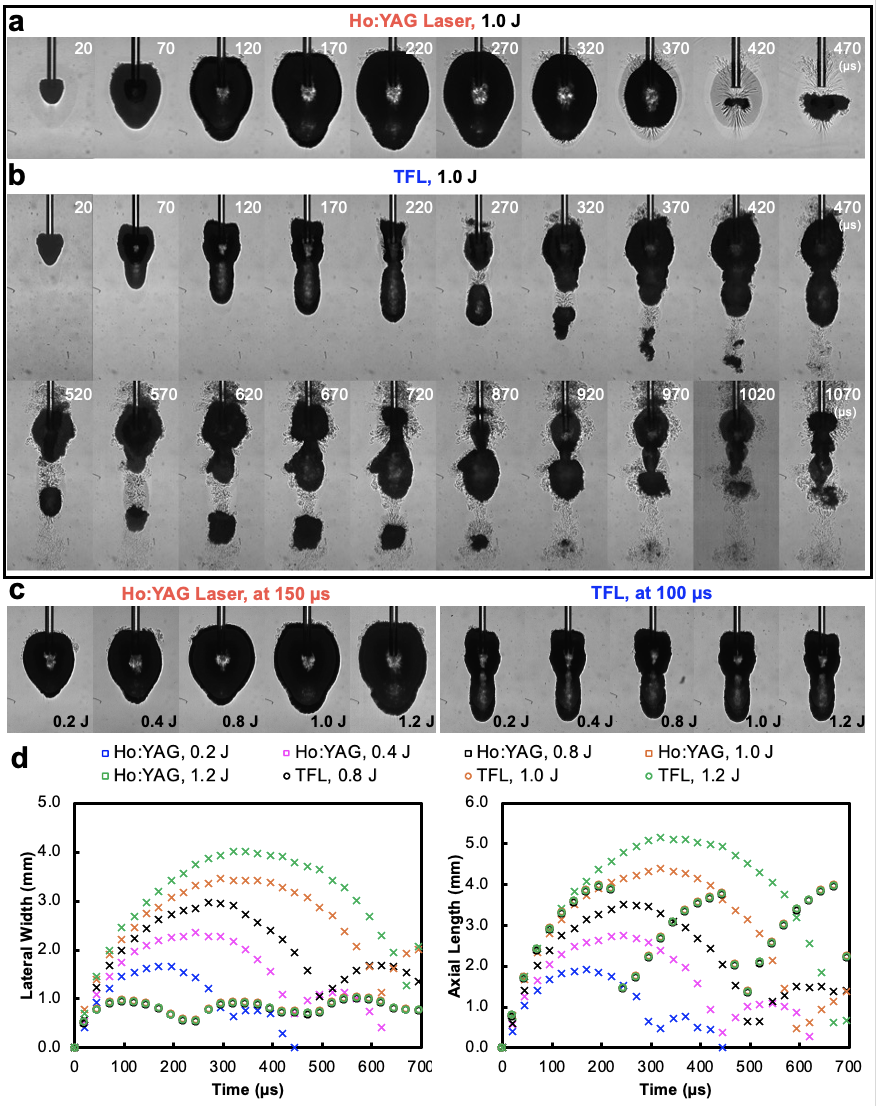

**Figure S3.** Bubble dynamics produced in a free field at 1.0 J by (a) Ho:YAG laser and (b) TFL. (c) Bubble morphology at 150 µs for Ho:YAG laser and 100 µs for TFL under varying pulse energy levels (0.2 J – 1.2 J) for both lasers. (d) Temporal evolution of bubble lateral width and axial length.
